# Epigenomic features related to microglia are associated with attenuated effect of APOE ε4 on alzheimer’s disease risk in humans

**DOI:** 10.1101/2020.09.28.317156

**Authors:** Yiyi Ma, Lei Yu, Marta Olah, Rebecca Smith, Stephanie R. Oatman, Mariet Allen, Ehsan Pishva, Bin Zhang, Vilas Menon, Nilüfer Ertekin-Taner, Katie Lunnon, David A. Bennett, Hans-Ulrich Klein, Philip L. De Jager

**Author notes:** Corresponding author: Philip L. De Jager, M.D., Ph.D., Director of the Center for Translational & Computational Neuroimmunology, Department of Neurology, Columbia University Medical Center, 630 West 168^th^ street, New York, NY 10032, USA.

## Abstract

**INTRODUCTION:** Not all *APOE* ε4 carriers who survive to advanced age develop Alzheimer’s disease (AD); factors attenuating the risk of ε4 on AD may exist.

**METHODS:** Guided by the top ε4-attenuating signals from methylome-wide association analyses (N=572, ε4+ and ε4-) of neurofibrillary tangles and neuritic plaques, we conducted a meta-analysis for pathological AD within the ε4+ subgroups (N=235) across four independent collections of brains. Cortical RNA-seq and microglial morphology measurements were used in functional analyses.

**RESULTS:** Three out of the four significant CpG dinucleotides were captured by one principle component (PC1), which interacts with ε4 on AD, and is associated with expression of innate immune genes and activated microglia. In ε4 carriers, reduction in each unit of PC1 attenuated the odds of AD by 58% (OR=2.39, 95%CI=[1.64,3.46], *P*=7.08×10^−6^).

**DISCUSSION:** An epigenomic factor associated with a reduced proportion of activated microglia appears to attenuate the risk of ε4 on AD.

## 1. Introduction

The *APOE* ε4 haplotype contributes the greatest common genetic risk for Alzheimer’s disease (AD)^1-4^. However, not all ε4 carriers develop AD. A longitudinal observational study reported that 9 out of the 141 ε4/ε4 individuals remained dementia-free after age 84^5^. Another meta-analysis of cross-sectional studies suggested that the ε4 effect on AD becomes weaker after age 70^1^. These results indicate that factors attenuating the genetic risk of ε4 on AD might exist. Such attenuators could be either genetic or environmental or both; here, we focused on DNA methylation features which might act as a modifiable mechanism on the human genome^6^.

Conducting a DNA methylation genome-wide association study (MWAS) of AD risk within the ε4+ subgroup can provide an unbiased search for those unknown signals protecting ε4 carriers from having AD. However, such an analysis is very limited by statistical power, as illustrated by our recent whole exome sequencing study with over 3,000 ε4+ subjects^7^. Therefore, we designed a three-stage approach (**Figure 1**). In stage I, we took advantage of an existing DNA methylation dataset^8^ generated from a random subset of participants in the Religious order (ROS) and Memory & Aging Projects (MAP). This includes both ε4+ and ε4-individuals with detailed quantitative measures of AD neuropathology to maximize power in performing an initial MWAS to prioritize a list of CpG dinucleotides which had the potential to be an ε4 attenuator. The resulting prioritized CpG dinucleotides fulfilled two criteria: (1) be associated with AD pathology in all subjects; and (2) reduce the effect of ε4 on AD susceptibility. The latter is determined by comparing the regression coefficients for the ε4 variable before and after adjusting for each CpG dinucleotide and focusing on those CpG dinucleotides which reduce the unadjusted ε4 regression coefficient. The prioritized list of CpG dinucleotides from stage I were then moved into stage II, where we checked their associations with AD in the ε4+ subgroup. We then assessed for evidence of replication in independent cohorts, and, to increase the statistical power, we conducted a meta-analysis in stage II to combine data from 4 independent cohorts and generate a final summary statistic. In stage III, we conducted validation analyses and explored the relevant functions. As a result, we have identified an epigenomic factor which might attenuate the ε4 effect on AD risk through changes in the transcriptome of the neocortex that relate to alterations in the proportion of activated microglia and their effect on the accumulation of Tau pathologies.

**Figure 1.**
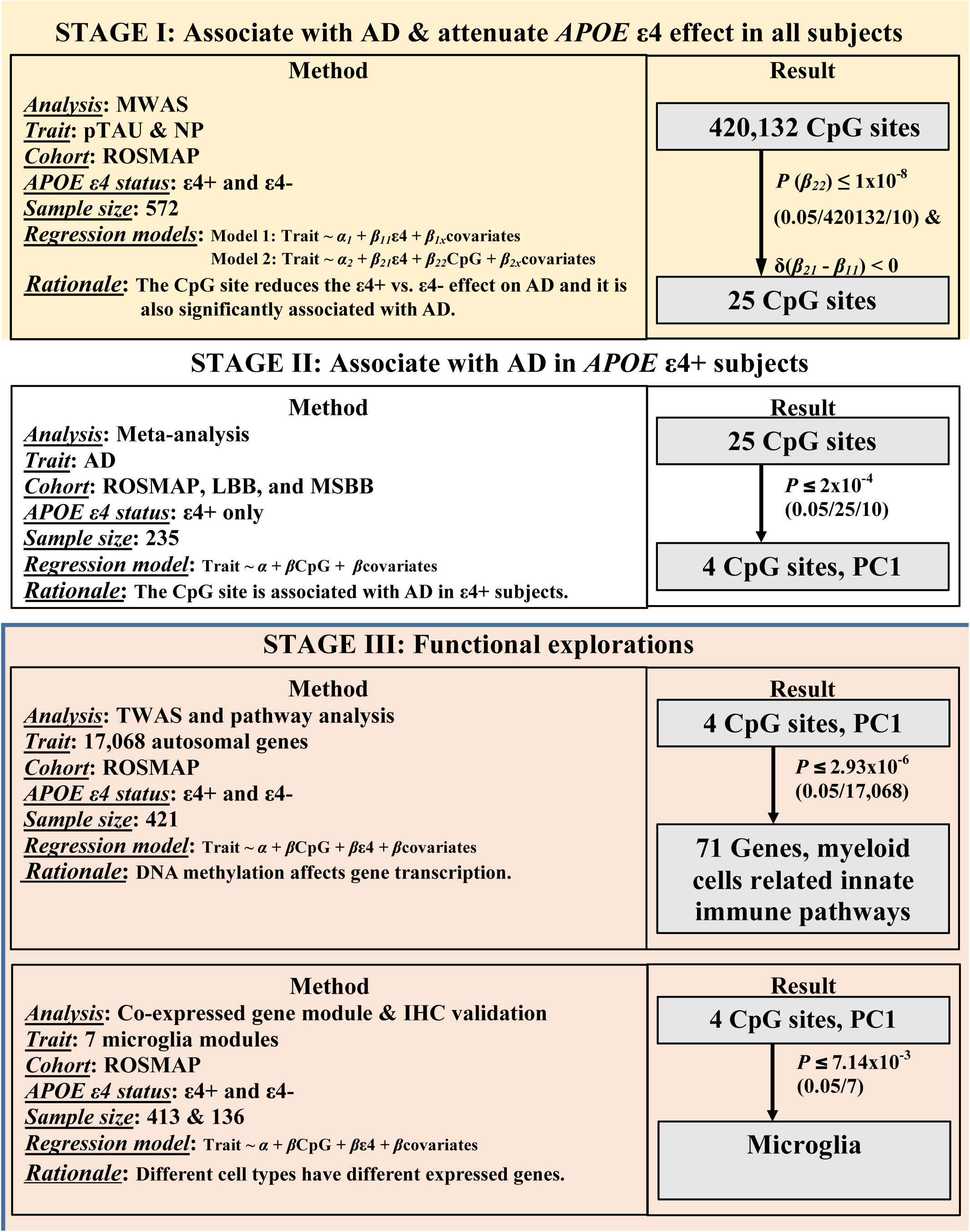
Flow chart of analysis plan and results. Abbreviations: MWAS, methylome-wide association analysis; NP, neuritic plaque; pTAU, abnormally phosphorylated Tau protein, AT8; TWAS, transcriptome-wide association analysis; IHC, immuno-histochemistry measurements.

## 2. Methods

### 2.1 Study samples and pathological AD measurements

We included whites from the studies of the Religious order Study (ROS) and the Rush Memory and Aging Project (MAP) (www.radc.rush.edu), the MRC London Neurodegenerative Disease Brain Bank (LBB), and the Mount Sinai Alzheimer’s Disease and Schizophrenia Brain Bank (MSBB) (**Supplementary Methods and Table S1**)^9-11^. ROS and MAP were jointly analyzed as ROSMAP with the adjustment of study variable (ROS and MAP). The LBB (GSE59685) and MSBB (GSE80970) included 68 (ε4+=36) and 129

(ε4+=41) individuals, respectively. Mayo Clinic Brain Bank (MAYO) provided DNA methylation and temporal cortex gene expression data on 45 patients with definite AD^12,13^, diagnosed neuropathologically according to NINCDS-ADRDA criteria^14^. Both temporal cortex and prefrontal cortex brain tissues were archived frozenly. The study was approved by IRB of each institute.

### 2.2 Pathological measurements

The pathological diagnosis of AD is based on the Braak score in LBB and MSBB and the NIA-Reagan score^15,16^ in ROSMAP, which relies on both neurofibrillary tangles (Braak) and neuritic plaques (NP). Details of common neuropathologic indices measured in the ROSMAP study were described before^17-20^ and in the **Supplementary Methods**.

### 2.3 Brain DNA methylation across studies

Details of DNA methylation measurements and data processing of the cortical samples of ROSMAP, LBB, and MSBB were described before^8-11^. Briefly, the genome-wide DNA methylation was measured by the Illumina 450K methylation array followed by QC and normalization^8,10,11,21^. As a result, β values for 420,132 CpG dinucleotides were included in the ROSMAP MWAS which yielded the 25 sites for subsequent meta-analysis across ROSMAP, LBB and MSBB. In MAYO, only cg05157625 is available out of the 4 top ones which was measured using the reduced representation bisulfite sequencing as described before^22^.

### 2.4 Brain gene expression in ROSMAP, MSBB, and MAYO

In ROSMAP, there were 421 subjects with both data of DNA methylation and RNA sequencing (RNA-seq) (Illumina) from their dorsolateral prefrontal cortex^9^ (Supplementary Methods). We included the 17,068 autosomal genes in the unit of normalized log2(cpm) into the transcriptome-wide association study (TWAS). A subset of these subjects (N=413) were previously used to derive the 47 cell-type relevant co-expressed gene module^23^. In MSBB, we downloaded the gene expressions of the transcriptome from Synapse platform (https://www.synapse.org/#!Synapse:syn7391833). Based on the genotype concordance check (Supplementary Methods), we included 50 subjects who have been profiled with both DNA methylation at prefrontal cortex and RNA-seq (Illumina) at BM44 region (closest to the prefrontal cortex).

In MAYO, we included 45 AD cases with both the DNA methylation data at cg05157625 and microarray-based gene expression data from their temporal cortex (Illumina)^12,13^.

### 2.5 Immunohistochemistry (IHC) measurements of cell types and microglia morphology in ROSMAP

A subset of ROSMAP subjects (N=57) with RNA-seq dataset were also profiled with the IHC stainings of markers of different cell types: neurons (NeuN), astrocytes (GFAP), microglia (IBA1), oligodendrocytes (Olig2) and endothelial cells (PECAM-1). The proportion of the microglia cell out of the total number of all different measured cell types were calculated^23^. Another subset (N=136) were evaluated for their microglia activation based on morphology changes: stage I (thin ramified processes), stage II (plump cytoplasm and thicker processes), and stage III (appearance of macrophages). The percentage or the square-root transformed proportion of the stage III activated microglia out of the sum of all three stages counts (PAM) were derived as before^24^. These measurements are pre-existing and independent from current study.

### 2.6 Statistical analysis

We used generalized linear regression model with the adjustments of age at death, sex, postmortem interval (if applicable), study (for ROSMAP dataset), technical variables, cell proportion^11^ (if applicable) and *APOE* ε4 carrying status (if necessary). We applied the Bonferroni-correction significance threshold. Considering the potential inflation by including the same subjects in stage I and II, we arbitrarily applied 10 times more stringent significance threshold in stage I (*P*≤1×10^−8^ (0.05/420,132/10)) and in stage II (*P*≤2×10^−4^ (0.05/25/10))). The correlations between pairs of the top 4 CpG dinucleotides were presented using R “ggcorrplot” package. The standardized β values (mean=0 and SD=1) of each of the top 4 CpG dinucleotides in ROSMAP were input to derive 4 principal components (PCs) in ROSMAP, which were further projected into LBB and MSBB using the R “factoextra” and “prcomp”

### 2.7 Pathway analysis

Top TWAS significant genes (*P*≤2.93×10^−6^ (0.05/17,068 autosomal genes)) were followed with pathway enrichment analysis using “STRINGdb” v10 against the functional Kyoto Encyclopedia of Genes and Genomes (KEGG) Pathway database (https://www.genome.jp/kegg/pathway.html)^25^ with FDR correction^26^.

### 2.8 Fetal brain Hi-C sequencing data downloaded from GEO

We downloaded (04/24/2020) the publically available Hi-C sequencing data of fetal brains^27^ (https://www.ncbi.nlm.nih.gov/geo/query/acc.cgi?acc=GSE77565) to interrogate the inter-chromosomal interactions. For each of the top 4 CpG dinucleotides, we extracted its genomic contacts across the 30,376 regions (100Kb resolution). Using the non-parametric Mann-Whitney-Wilcoxon test, we compared whether the rank of the normalized contact frequency is different between the two types of regions (TWAS or non-TWAS regions).

## 3 Results

### 1.1. Demographic characteristics by APOE ε4

In ROSMAP, compared to *ε4*-, *ε4*+ participants were younger at the time of death (*P*=0.02), more likely to have pathological AD (*P*=2.48×10^−9^), and had more abnormally phosphorylated tangles (pTAU) (*P*=2.61×10^−8^) and neuritic amyloid plaques (NP) (*P*=5.6×10^− 11^) (**Table 1**). In the MRC London Neurodegenerative Disease Brain Bank (LBB) and the Mount Sinai Alzheimer’s Disease and Schizophrenia Brain Bank (MSBB) datasets, *ε4*+ subjects were also more likely to have AD (*P*<0.005) than *ε4*-subjects.

**Table 1.**
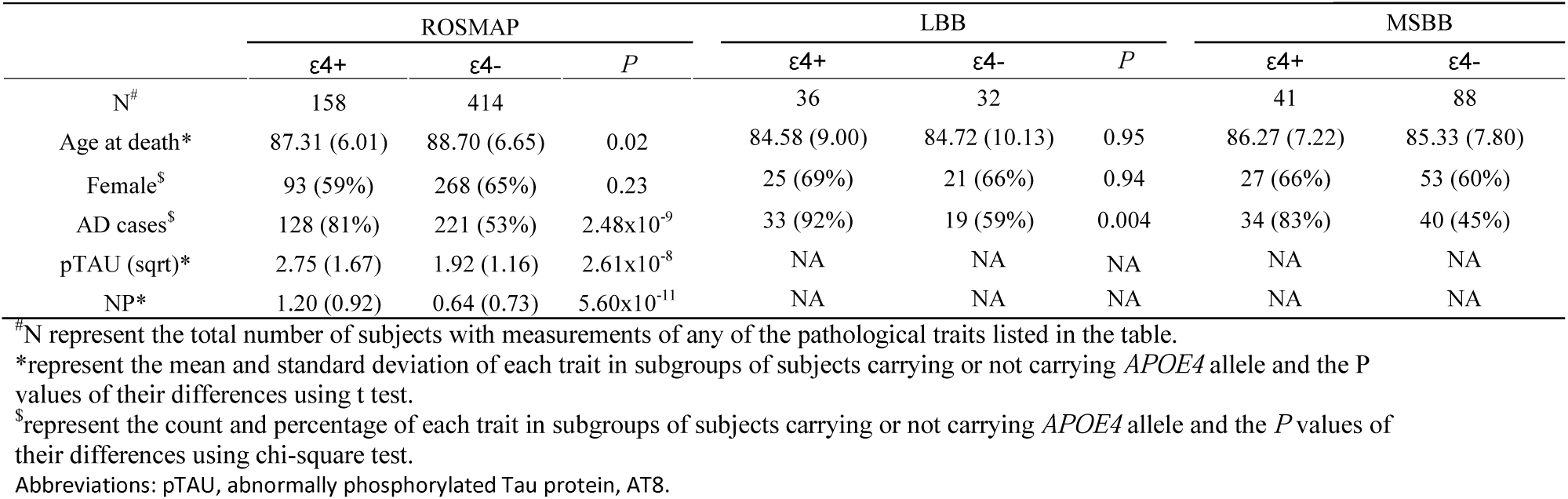
Demographics of subjects carrying or not carrying *APOE ε4* allele across ROSMAP, LBB and MSBB

### 1.2. Identification of CpG dinucleotides attenuating the effect of APOE ε4 on AD

#### 3.2.1 Discovery of CpG dinucleotides

In our stage I analysis with all ROSMAP subjects (**Figure 2A & Table S2)**, 25 CpG dinucleotides (1) were significantly associated with either pTAU or NP (*P*≤1×10^−8^) and (2) had potential to attenuate the effect of ε4 on pTAU or NP because the regression coefficients of *APOE* ε4, after adjusting for the candidate CpG dinucleotide, were smaller than the unadjusted ones. These 25 CpG were then evaluated in stage II, where we conducted a meta-analysis across 235 *ε4*+ individuals assembled from four sample collections: ROS, MAP, LBB and MSBB. We found 4 CpG associated with a pathologic diagnosis of AD (meta-*P*≤2×10^−4^): cg08706567 (*MPL*), cg26884773 (*TOMM20*), cg12307200 (*LPP* and *TPRG1*), and cg05157625 (*RIN3*) (**Figure 2B**). All of these 4 CpGs have stronger effects on AD susceptibility in *ε4*+ than *ε4*-, which is consistently observed across cohorts (**Figure 2C**,**S1**). For the cg08706567 and cg26884773, the LBB *ε4*+ dataset was not included into the meta-analysis since it has the problem of infinite maximum likelihood estimates (*P*=1)^28^, and the results using the penalized generalized regression model yielded significant meta-P (*P*≤0.01) including only the LBB and MSBB (**Table S3**). Thus, the independent cohorts offer evidence of replication.

**Figure 2.**
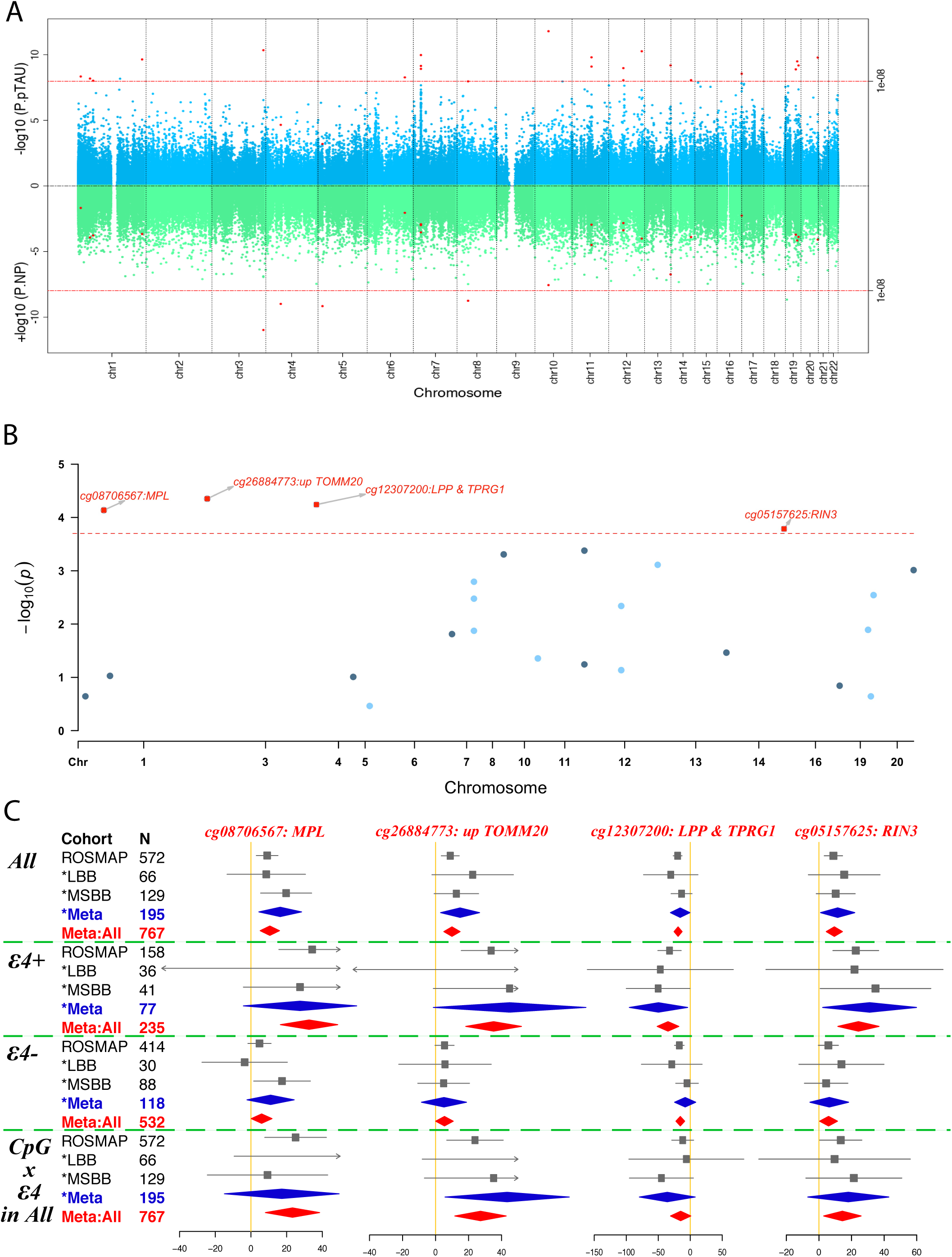
Identification of the top 4 CpG dinucleotides. (A) Miami plot shows the results of the methylation genome-wide association study (MWAS) on pTAU (upper panel in blue) and NP (lower panel in green) in ROSMAP (N=572). The Y axis show the -log10 transformed *P* value of each of the genome-wide CpG dinucleotides which are shown in dots and ordered according to their genomic coordinates on the shared X axis. The red dashed line represents the significance threshold (*P*=1E-8=0.05/420132/10) and those CpG dinucleotides passing the genome-wide significance threshold for either pTAU or NP are shown in red dots. (B) Candidate manhattan plot shows the meta-analyzed associations of the above selected 25 CpG dinucleotides with pathological AD in _ε_4+ subjects across ROSMAP, LBB, and MSBB cohorts (N=235). Y axis represent the -log10 transformed meta analyzed *P* values of the regression coefficient estimates of the logistic regression model, in which the outcome variable is the pathological diagnosis of Alzheimer’s disease (AD) (no=0 and yes=1), the exposure variable is the methylation status of each CpG dinucleotides (0 to 1) and the covariates include age at death, postmortem interval, sex, and study, ethnicity principle components, methylation experiment batches in ROSMAP, and cell proportion in LBB and MSBB. The horizontal red dashed line represents the Bonferroni corrected *P* value threshold of (0.05/(25*10) = 2×10^−4^) and the top 4 significant CpG dinucleotides are represented in red squares with the annotation of their closest genes. (C) Forest plots for the top 4 CpG dinucleotides across different cohorts for their associations with AD within: 1) all the subjects; 2) subjects carrying; or 3) not carrying the *APOE* _ε_4 allele; and 4) the interaction test between the methylation level and *APOE* _ε_4 allele carrying status within all the subjects. The filled square and horizontal line for each population or the filled summary diamonds (blue for the meta-analysis across the replication cohorts while red for the joint meta-analysis across all the cohorts) denote the estimated regression coefficient (BETA) and its 95% CI per unit increase in the methylation level of each CpG dinucleotide or the interaction term of the methylation times the _ε_4-carrying status (yes=1 and no=0) for the binary outcome variable of the pathological diagnosis of AD (no=0 and yes=1) with the adjustment of the covariates of age at death, postmortem interval, sex, study, ethnicity principle components, methylation experiment batches in ROSMAP, and cell proportion in LBB and MSBB. The arrows indicate the estimates are out of the boundaries. Abbreviations: NP, neuritic plaque; pTAU, abnormally phosphorylated Tau protein, AT8.

#### 3.2.2 Pathological associations of the 4 CpGs

Aside from its strong associations with pTAU and NP, cg12307200 had weaker associations with diffuse plaques (BETA=-3.15, SE=1.03, *P*=2.34×10^−3^), cerebral amyloid angiopathy (BETA=-3.36, SE=1.19, *P*=5.07×10^−3^), and arteriosclerosis (BETA=-3.47, SE=1.28, *P*=6.73×10^−3^); and it had no association with Lewy Bodies, hippocampal sclerosis, or TDP-43 proteinopathy (*P*>0.05). The other 3 CpG dinucleotides had weaker associations with NP (BETA=[3.51,4.07], SE=[0.94,0.5], *P*=[1.15×10^−4^,2.14×10^−4^]) than pTAU (BETA=[8.88,9.81], SE=[1.49,1.66], *P*=[1.33×10^−5^,1.64×10^−4^]), nominal associations with TDP-43 proteinopathy (BETA=[2.28,3.24], SE=[1.36,1.51], *P*=[0.03,0.09]), and no associations with other neuropathologic indices. Thus, the top 4 CpG dinucleotides were primarily associated with Tau-related pathologies (Table S2).

#### 1.3. Derivation of the methylation PC based on the identified top 4 CpGs

All of the subjects have hypermethylation (β≥0.5) at cg12307200 and hypomethylation (β<0.5) at the other 3 CpGs. While they are located on different chromosomes, the methylation level of these 3 CpGs were correlated (*r*≥0.4) (**Figure 3A**,**S2**). Because such patterns were consistently observed across all the cohorts, we derived a methylation PC1 that captures the effect of those 3 highly-correlated CpGs in a single measure. We developed it using ROSMAP data and then projected it to the samples from LBB and MSBB using eigenvectors. As expected, the projected PC1 in LBB and MSBB captured the 3 CpG dinucleotides in the same way as in ROSMAP (**Figure 3B**).

**Figure 3.**
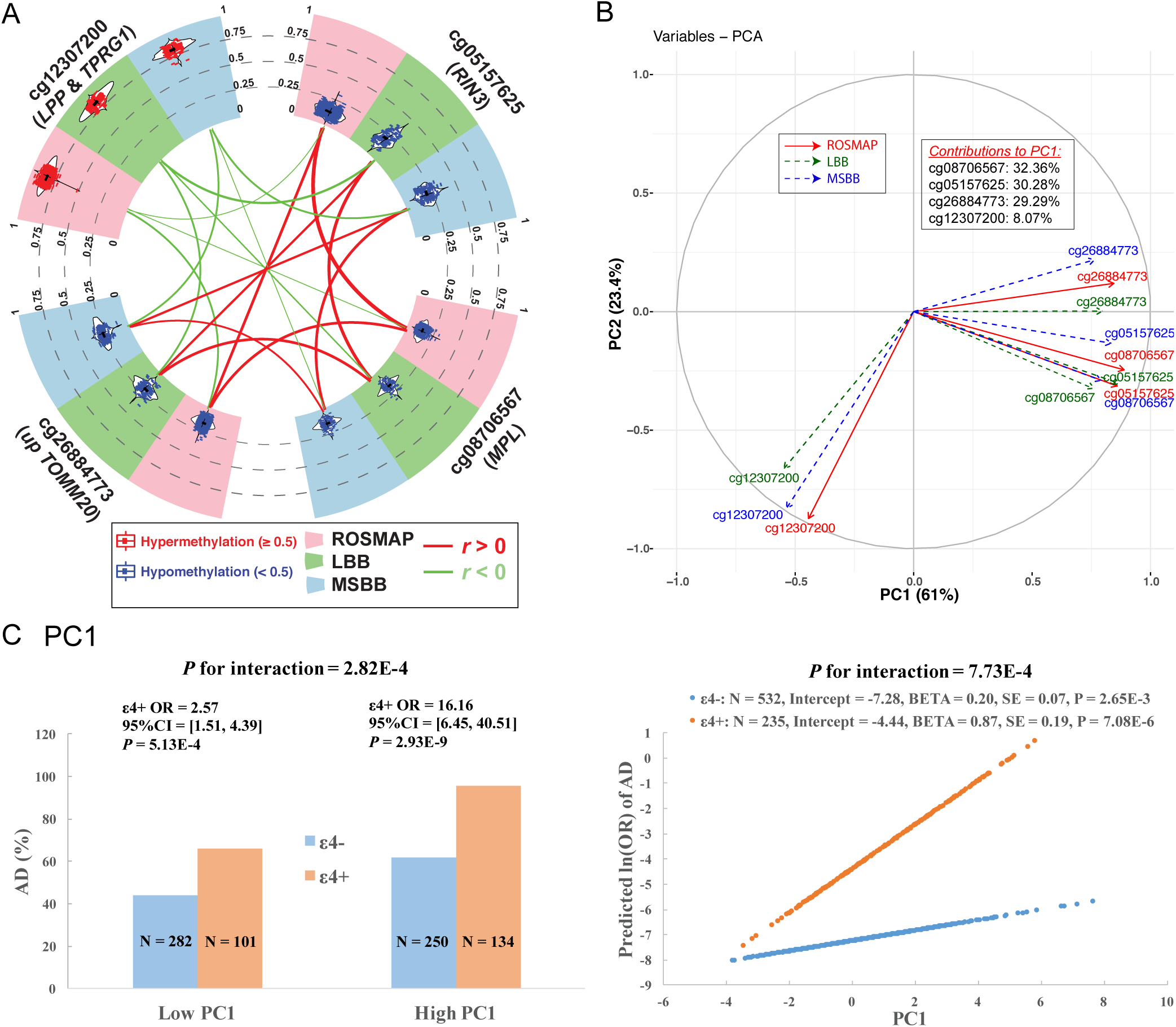
Consistent correlations among the 4 CpG dinucleotides across cohorts and the derivation of the PC1. (A) Circos plot shows the consistent distributions of each of the top 4 CpG dinucleotides and their mutual correlations across ROSMAP (pink sector), LBB (green sector) and MSBB (blue sector). In the outer layer, the distribution of each CpG dinucleotide within each cohort is shown in a violin plot where each dot represents a subject and the black horizontal and vertical lines denote the mean and standard deviation. Red dots represent those subjects with hypermethylation level (_≥_0.5) while blue ones represent those subjects with hypomethylation level (<0.5). In the inner layer, the correlations between each pair of 2 CpG dinucleotides within each cohort were represented by their connected lines, where the color denote the direction (red for *r*>0 and green for *r*<0) and the thickness denote the strength of their correlations (thicker lines represent stronger correlations). (B) Variance plot of the PC1 vs. PC2 and their contributions to each of the top 4 CpG dinucleotides across ROSMAP (red solid arrow), LBB (blue dashed arrow), and MSBB (blue dashed arrow). (C) Interaction effect on AD between *APOE* _ε_4 and PC1. The subgroup of _ε_4+ and _ε_4- are represented as blue and orange. Interaction with categorical variables are shown on the left panel where the continuous variable of PC1 were transformed to the binary variable based on the median value. The Y axis represent the percentage of AD cases across 4 subgroups of subjects. The right panel show the interactions with the untransformed continuous PC1. the Y axis represent the predicted ln(OR) of AD calculated based on the statistics within _ε_4+ and _ε_4-subgroup using the logistic regression model with binary AD status (case=1 and control=0) as the outcome variable adjusting the covariates of age at death, sex, study, postmortem interval, methylation experimental batches and two major ethnic principles. Abbreviations: OR, odds ratio.

### 1.4. Interaction between APOE ε4 and the top CpG dinucleotides on AD risk

Here, we evaluated whether our epigenomic factors interacted with *APOE* ε4 to modulate AD risk and compared their stratified effects on AD by ε4. The cg12307200 site did not display significant evidence of interaction, but the 3 correlated CpG dinucleotides captured by PC1 had nominal significance for ε4 interaction (range of meta-*P* for interaction=[6.1×10^− 4^,0.01]) (**Figure 3C**). This led us to pursue a more comprehensive interaction analysis of PC1 both categorically and continuously. For the categorical interaction, the original continuous PC1 was transformed to a binary variable based on its median value. For the continuous interaction, the non-scaled continuous value of the PC1 was used. The results were similar for both the non-scaled and scaled values (**Table S4**), ruling out the potential influences of outliers on the continuous analysis. Both the categorical (meta-*P*=2.8×10^−4^) and continuous (meta-*P*=7.7×10^−4^) interaction tests were significant. In the categorical analysis, the effect of ε4 on AD was smaller in the subjects with low PC1 (meta-OR=2.57, 95% CI=[1.51,4.39], *P*=5.13×10^−4^) than the subjects with high PC1 (meta-OR=16.16, 95% CI=[6.45,40.51], *P*=2.93×10^−9^). In the continuous analysis, the effect of PC1 on AD risk was greater in ε4+ (meta-ln(OR)=0.87, meta-*P*=7.08×10^−6^, meta-N=235) than in the ε4- (meta-ln(OR)=0.20, *P*=2.65×10^−3^, meta-N=532). In other words, reduction of one unit in PC1 was associated with a 58% ((1-1/exp(0.87))) attenuation in AD risk in ε4+ carriers. Thus, one could potentially influence the magnitude of ε4 risk by modulating PC1, an epigenomic measure that captures the effect of multiple loci.

### 1.5. Functional exploration neocortical ROSMAP transcriptomes, replications in other independent studies and validations by Hi-C sequencing data of fetal brains

To understand the functional consequences of our 4 CpG dinucleotides, we then conducted a Transcriptome-wide Association Study (TWAS) in the 421 ROSMAP participants who have both DNA methylation and RNA sequence (RNA-seq) data from the same cortical region. There were 71 genes which displayed significant (*P*≤2.93×10^−6^ (0.05/17,068 autosomal genes)) associations with either PC1 (70 genes) or cg12307200 (3 genes) (**Figure 4A&B, Table S5**). Except for *RIN3* where there was a cis-effect of the DNA methylation on the gene expression, all the other associations are driven by the trans-effects because the TWAS target genes are far from the 4 CpG dinucleotides: they are either on different chromosomes or >5 Mb apart for those on the same chromosome.

**Figure 4.**
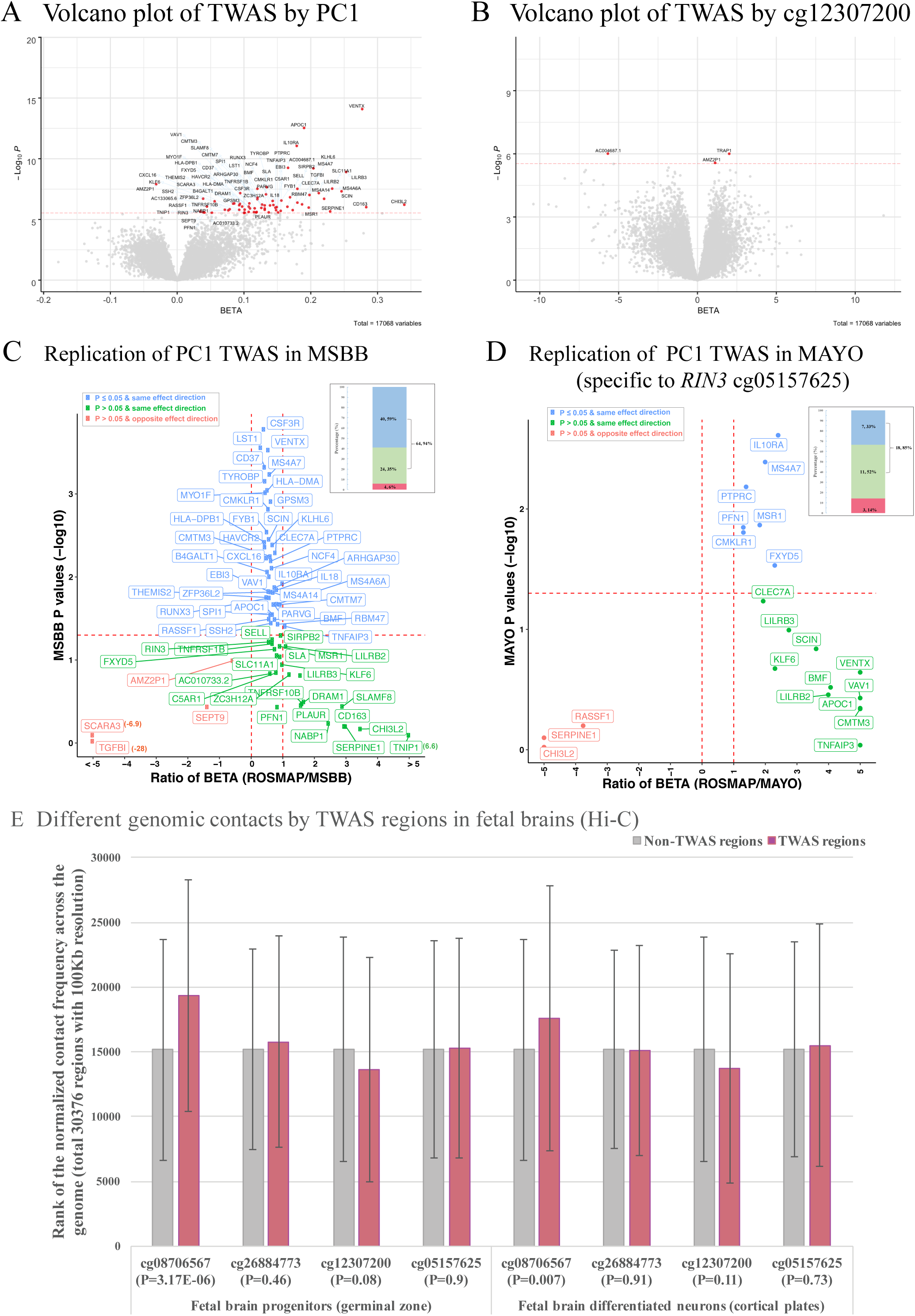
Discovery and replication of TWAS results of the PC1 and cg12307200. (A) and (B) Volcano plot shows the TWAS results of PC1 and cg12307200. The X axis shows the BETA and the Y axis shows the -log10 transformed P values for the exposure variable of PC1 or cg12307200 on the mRNA expression levels of each gene. The 71 unique genes (70 for PC1 and 3 for cg12307200) passed the Bonferroni corrected significance threshold of *P* _≤_ 2.93×10^−6^ (0.05/17,068 autosomal genes) are shown in red dots with their gene name connected through blue arrows. (C) and (D) Replication of the PC1 TWAS results in MSBB and MAYO. ROSMAP has identified 70 TWAS genes for PC1 and 68 of them are available in MSBB. There are 25 (out of 70) genes are significant for cg05157625 (the only CpG dinucleotide available in MAYO) and 21 of them are available in MAYO. The scatter plot shows the comparison of each gene between ROSMAP and MSBB (or MAYO) where the X axis represents the ratio of the regression coefficient obtained in the two cohorts (ROSMAP over MSBB (or MAYO) and the red dashed vertical lines for the ratio=0 or 1) and the Y axis shows the *P* values in MSBB (or MAYO) (red dashed horizontal line for the *P*=0.05). The blue dots are those genes with nominal significance (*P*_≤_0.05) in MSBB (or MAYO) and with the same effect direction in both ROSMAP and MSBB (or MAYO), the green dots are those non-significant genes (*P*>0.05) in MSBB (or MAYO) but with the same effect direction, and the red dots are those non-significant genes in MSBB (or MAYO) and also with the opposite effect direction. The number and their percentage of these three groups of genes are presented in the bar plot (upper right corner) with the same color codings. (E) Different genomic contacts by TWAS regions in the fetal brains analyzed with the published Hi-C sequencing data (Won et al., 2016) downloaded from GEO (GSE77565). Silver and rose gold bars represent those regions outside and within the 70 significant TWAS genes in ROSMAP and the *P* value for the differences between these two groups for each CpG dinucleotide are shown in X axis.

We then attempted to replicate our TWAS findings. In MSBB (50 subjects), the effect of cg12307200 was not replicated. But the majority of the PC1 TWAS genes were replicated. Out of the total of 68 genes also available in MSBB, 94% have the same effect direction as in ROSMAP, and 59% also showed significance (*P*<0.05) (**Figure 4C**). In MAYO (45 AD cases), only one of the three PC1 CpGs (cg05157625) was available. Out of the 21 TWAS genes significant with both PC1 and cg05157625 in ROSMAP, 85% have the same effect direction in MAYO as in ROSMAP (**Figure 4D**). 4 genes are significant in both MAYO and MSBB in relation to PC1, despite the small sample sizes.

To further explore the biological grounding of our observation, we accessed the publically available Hi-C sequencing data from human fetal brain tissue which captures the 3-dimensional architecture of human cortex, with the caveat that this profiles a stage of brain development. Despite this important limitation, we found evidence that those genomic regions containing the 71 TWAS genes identified in ROSMAP have more contacts with the region covering one of the three PC1 CpGs, cg08706567, than you would expect by chance (*P*<0.01) (**Figure 4E**). This suggests that, even at very early stages of brain development, the 71 TWAS genes are physically interacting with at least one element of PC1; cg08706567 may play an important role as a regulator locus whose impact continues into advancing as it influences the impact of ε4.

### 1.6. The TWAS results implicate microglial activation in the effect on ε4

#### 3.6.1 Pathway analysis of TWAS results suggested the involvement of the myeloid cells

These 71 TWAS significant genes displayed enrichment for 20 KEGG functional pathways (FDR≤0.05, hits>1 and hits%≥3%) (**Figure 5A**), which were related to multiple aspects of immunity and particularly myeloid cell function, such as osteoclast differentiation (hits=4%, FDR-*P*=5.5×10^−5^), phagosome (hits=4%, FDR-*P*=5.5×10^−5^), tuberculosis (hits=3%, FDR-*P*=5.5×10^−5^), Leishmaniasis (hits=4%, FDR-*P*=1.3×10^−3^), and antigen processing and presentation (hits=3%, FDR-*P*=0.01).

**Figure 5.**
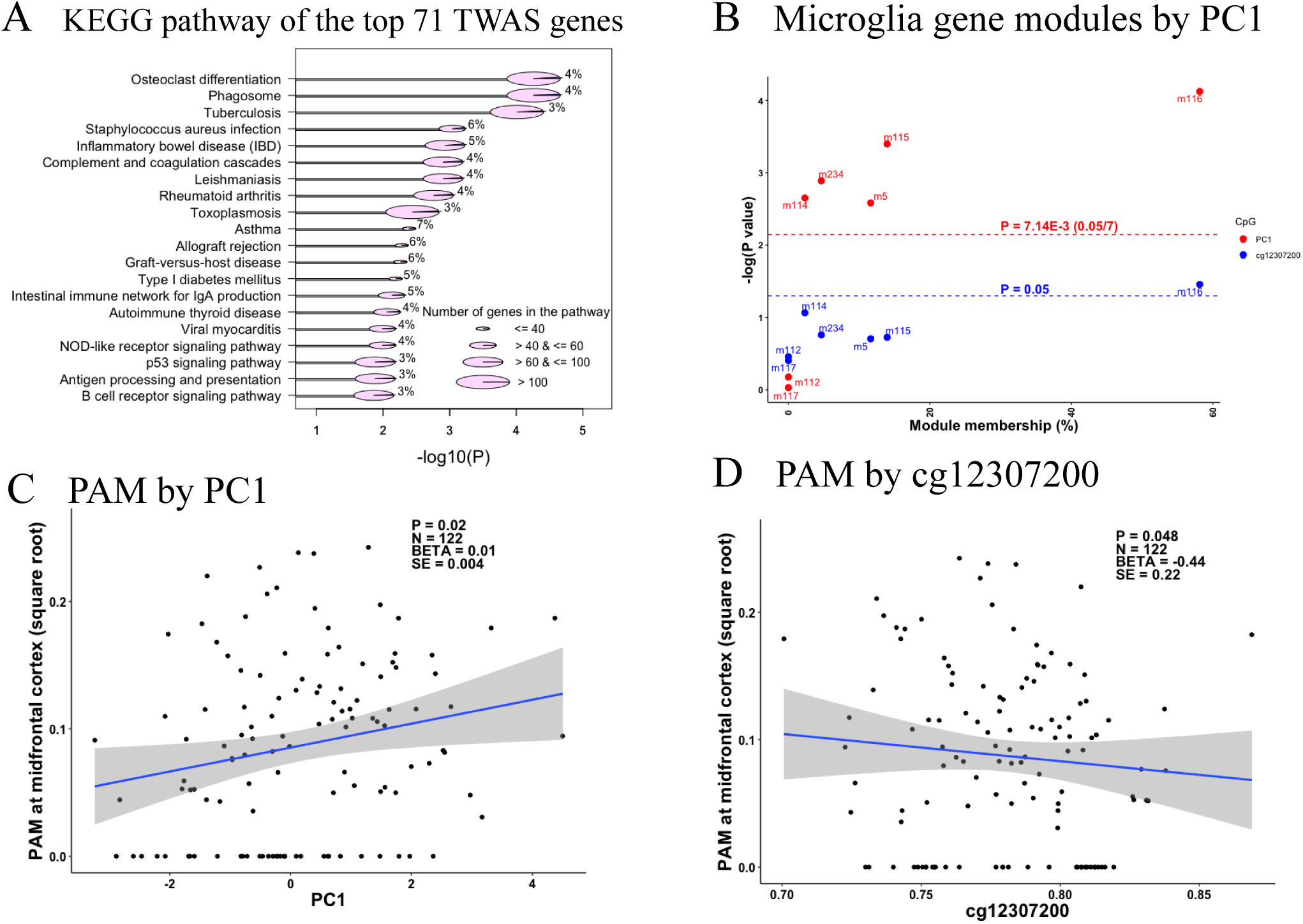
Microglia relevance to PC1 and cg12307200. (A) Plot of the KEGG pathway analysis of the top 71 TWAS genes which have significant associations with the PC1 and cg12307200. Only those pathways with FDR adjusted *P* value _≤_ 0.05 and the number (percentage) of the hit genes > 1 (_≥_ 3%) were shown in the plot. Y axis list the name of these KEGG functional pathways and X axis shows their corresponding –log10 transformed FDR adjusted *P* values. The size of the pie represents the number of the member genes of each functional pathway, which are categorized into four types ranging from the smallest to the largest containing _≤_40, >40 & _≤_60, >60 & _≤_100, and >100 member genes. Each pie was split into 2 slices with the area of the blue slice represents the percentage of the hit genes out of the total member genes for each pathway and their number are shown for each pathway. (B) Scatter plot of the associations with the methylation factors (PC1 in red and cg12307200 in blue) and expression levels of the 7 reported gene modules for microglia. Y axis represent the –log10 transformation of the *P* values of the associations between the exposure variables of PC1 or cg12307200 on the outcome variable of the expression values of each gene module, and the X axis represent the module membership, which is defined as the percentage of the module member genes out of our identified 43 gene list (28 genes are not mapped to any of the reported gene modules). The horizontal red and blue dashed lines represent the Bonferroni corrected significance threshold of *P*_≤_7.14×10^−3^ (0.05/7 gene modules) and the nominal significance threshold of *P*_≤_0.05. (C) and (D) show the scatter plot of the associations between proportion of activated microglia (PAM) at midfrontal cortex as Y axis and the methylation factors (PC1 and cg12307200) as X axis. Each dot is one observation and the regression line and its 95% CI are represented by the blue line and grey area. The estimated statistics are shown on the upper right corner of the plot. The BETA, SE, and P represent the estimated regression coefficient and its standard error, P value of the exposure variable of PC1 or cg12307200 on the outcome of PAM. The N represent the sample size within the analysis. For all the analysis, we used the generalized linear regression model with the adjustments of the age at death, sex, postmortem interval, study, *APOE* _ε_4 binary status, two major ethnic principles and methylation experimental batches.

#### 3.6.2 Association with co-expressed gene modules of microglia

We further conducted a complementary association analysis with the 7 modules of co-expressed genes previously described as being enriched for microglial genes ^23,29^: m5, m113, m114, m115, m116, m112, and m117. PC1 was associated with the expression of 5 of these 7 microglia modules (*P*≤7.14×10^−3^ (0.05/7 microglia modules)) (**Figure 5B)**, but not other non-microglia modules (**Table S6**). Further, almost all (91%) of the 71 TWAS genes were found in the 5 significant microglia modules, and 58% belonged to m116, the most microglial enriched module; we previously reported^23^ this module as being related to microglial aging^29^. It also contains some key AD genes such as *TREM2* and its binding partner *TYROBP*. m5 has been associated the burden of Tau pathology and the proportion of activated microglia (PAM), as defined using data from immunohistochemistry of brain sections and a standard neuropathologic scale^24,29^. Our findings do not appear to be driven by changes in microglial cell counts since neither PC1 nor cg12307200 was associated with the microglia cell counts estimated by either the immunohistochemistry staining for IBA1 protein or its mRNA expression levels (**Table S7**). The results remain significant when we account for the proportion of microglial cells. Overall, our 4 epigenomic factors appeared to be related to microglial transcriptional programs captured by the modules of co-expressed genes defined in neocortical data, and the state of the microglia may thus be an important factor in the modulation of the ε4 effect.

#### 3.6.3 Validation using a histology-derived variable of microglia activation

As noted above, the m5 module was associated with PAM, a trait derived from the morphological characterization of microglia in histological sections that we found to be associated with cognitive decline, AD pathologies, and AD dementia^24^. PAM is simply the proportion of microglia that have an activated stage III morphology, and this trait was available in 122 ROSMAP participants that also have DNA methylation data from the same frontal cortex region. PC1 was positively (*P*=0.02) and cg12307200 was negatively associated with PAM in the midfrontal cortex (*P*=0.05) (**Figure 5C,D**). However, the association is modest, and PC1 was not fully explained by PAM. That is, PAM and PC1 capture somewhat different aspects of microglial function that may be partially related to one another. To be thorough, we also evaluated PAM measures in three other brain regions in secondary analyses, and we found that PAM in inferior temporal cortex, posterior putamen and ventral medial caudate also displayed association with PC1 and cg12307200 (**Figure S2**), suggesting that PC1 and cg12307200 derived from cortical tissue DNA methylation data capture an aspect of microglial state in multiple brain regions, not just the frontal cortex.

## 4. Discussion

Our study explored the epigenome of the human brain for CpG dinucleotides that attenuate the impact of the strongest but not deterministic^5^ genetic risk of AD in the general population: the *APOE* ε4 haplotype. Our staged approach identified 4 primarily Tau-related CpGs and three of them were captured by one principal component (PC1). PC1 is positively associated with the proportion of activated microglia and had a significant interaction with ε4 showing stronger effect in ε4+ than ε4-subjects. Each unit reduction of PC1 in ε4 carriers attenuated AD risk by 58%, suggesting that we may have found a meaningful ε4 attenuator. Further work is now needed to further characterize PC1 to evaluate whether its effect could potentially be mimicked by a therapeutic agent.

We took advantage of the large and varied sets of molecular and histological data available in the same brain region of ROSMAP participants from which the DNA methylation profile were obtained to begin to characterize the biology captured by our 4 CpG dinucleotides. Our initial unbiased analysis clearly pointed towards the innate immune system and particularly myeloid cells, leading us to focus our attention on this cell type as we dissected the effect of the 4 CpGs. Namely, they seem to involve effects on two modules of co-expressed neocortical genes: module 116 which relates to microglial aging and module 5 which is associated with proportion of activated microglia and contributes to the accumulation of Tau pathology^29^. We previously reported that the proportion of the activated microglia was an independent risk factor for AD that had an effect size comparable to that of ε4 on AD^24^. Our current findings with the same dataset therefore refine our understanding of the relationship between activated microglia and the ε4 allele: while their effects on risk may be largely independent, they may interact to some degree.

Our findings that the top 4 CpG dinucleotides were primarily associated with Tau-related pathologies are in line with previous genetic studies revealing the link between *APOE* and *MAPT*, the gene encoding the Tau protein. A *MAPT* variant enriched in *MAPT* H2 carriers conferred greater protection from AD in *APOE* ε4-individuals^7,30^. A recent *MAPT* haplotype stratified GWAS identified a variant enriched in *APOE* ε3-carriers and protective in *MAPT* H1H1 individuals^31^. These findings alongside ours suggest the presence of either genetic or epigenetic factors which behave in a differential manner either on the *APOE* or *MAPT* genetic background to modify AD risk. Further, these context-specific factors could establish a link between *APOE* (major risk factor for AD) and *MAPT* (major risk factor for tauopathies).

Our study has several limitations. The sample size is limited even though we have assembled the largest ε4+ study with both DNA methylation and neuropathology data. Having low power is a common issue for all ε4+ subgroup studies^7^. Trying to circumvent this issue, we utilized a staged approach followed by RNA- and histology-based validation. The inclusion of the same ROSMAP subjects in both stage I and stage II may inflate our results, although we use different traits in each stage and impose a 10 fold more stringent significance threshold to address, in part, this issue. Finally, our cross-sectional design cannot to determine the causality, which is a general limitation of all postmortem autopsy studies for which we can only have one time point.

## 5. Conclusions

We reported that the deleterious effect of the strongest genetic risk (*APOE* ε4) for AD may be attenuated by an epigenomic factor which could work, at least in part, through alterations in the relative proportion of activated microglia and, subsequently, on the accumulation of Tau pathology. Further mechanistic studies are necessary to validate our results and demonstrate the sequence of events outlined in the hypothesis suggested by our current findings.

## SUPPLEMENTAL MATERIAL CONTENTS

### SUPPLEMENTAL METHODS (Text)

#### SUPPLEMENTAL TABLES

**Table S1**. Population demographics included in each analysis across all studies.

**Table S2**. Summary statistics of the 25 CpG dinucleotides for their associations with the 11 common neuropathologies in all subjects of ROSMAP, and meta-interactions with APOE ε4 on pathological AD in all subjects across ROSMAP, LBB, and MSBB, and meta-associations with pathological AD in subgroups of subjects carrying or not carrying ε4 allele across ROSMAP, LBB, and MSBB (in excel spreadsheet).

**Table S3**. Meta-analysis of cg08706567 and cg26884773 using penalized generalized regression model.

**Table S4**. Interaction and association tests with scaled continuous variables.

**Table S5**. Summary statistics of the top 71 genes from the transcriptome wide association for the PC1 and cg12370200 in ROSMAP with the adjustment of *APOE* ε4 status MSBB (in excel spreadsheet).

**Table S6**. Summary statistics of the association between the 47 gene modules and the PC1 and cg12370200 in ROSMAP with the adjustment of *APOE* ε4 status MSBB (in excel spreadsheet). **Table S7**. Associations of PC1 and cg12307200 with microglia cell type proportion in subset of subjects with immuno-histochemistry measurements (N=57).

#### SUPPLEMENTAL FIGURES

**Figure S1**. Forest plots of the association between the pathological diagnosis of Alzheimer’s disease (AD) and methylation level at the 4 top CpG dinucleotides.

**Figure S2**. Distributions of the scaled values of each of the top 4 CpG dinucleotides within each cohort before the derivation of PC1 and their pairwise correlations.

**Figure S3**. Associations of the proportion of activated microglia (PAM) with the methylation PC1 and cg12307200.

## DATA AVAILABILITY

All the data and analysis output are available via the AD Knowledge Portal (https://adknowledgeportal.synapse.org). The AD Knowledge Portal is a platform for accessing data, analyses, and tools generated by the Accelerating Medicines Partnership (AMP-AD) Target Discovery Program and other National Institute on Aging (NIA)-supported programs to enable open-science practices and accelerate translational learning. The data, analyses and tools are shared early in the research cycle without a publication embargo on secondary use. Data is available for general research use according to the following requirements for data access and data attribution (https://adknowledgeportal.synapse.org/DataAccess/Instructions). The link to the data and analysis output for this manuscript is https://www.synapse.org/#!Synapse:syn22240706.

## ACKNOWLEDGMENTS

<u>ROSMAP</u>: We are grateful to the participants in the Religious Order Study, the Memory and Aging Project. This work is supported by the US National Institutes of Health [U01AG61356, R01 AG043617, R01 AG042210, R01 AG036042, R01 AG036836, R01 AG032990, R01 AG18023, RC2 AG036547, P50 AG016574, U01 ES017155, KL2 RR024151, K25 AG04190601, R01 AG30146, P30 AG10161, R01 AG17917, R01 AG15819, K08 AG034290, P30 AG10161 and R01 AG11101. Ma Y. is supported by the 2019-AARF-644521.

<u>MSBB</u>: Brain banking and neuropathology assessments for the Mount Sinai cohort was supported by US National Institutes of Health grants AG02219, AG05138, U01 AG046170, RF1 AG057440 and MH064673, and the Department of Veterans Affairs VISN3 MIRECC. Replication work in Boston was supported by US National Institutes of Health grants: R01 AG036042, R01AG036836, R01 AG17917, R01 AG15819, R01 AG032990, R01 AG18023, RC2 AG036547, P30 AG10161, P50 AG016574, U01 ES017155, KL2 RR024151 and K25 AG041906-01.

<u>LBB:</u> Brain banking and neuropathology assessments for the MRC London Neurodegenerative Diseases Brain Bank, which was supported by the Medical Research Council (UK), and Brains for Dementia Research (Alzheimer Brain Bank, UK).

<u>MAYO</u>: Study data^13^ were provided by the following sources: The Mayo Clinic Alzheimers Disease Genetic Studies, led by Dr. Nilüfer Ertekin-Taner and Dr. Steven G. Younkin, Mayo Clinic, Jacksonville, FL using samples from the Mayo Clinic Study of Aging, the Mayo Clinic Alzheimers Disease Research Center, and the Mayo Clinic Brain Bank. Data collection was supported through funding by NIA grants P50 AG016574, R01 AG032990, U01 AG046139, R01 AG018023, U01 AG006576, U01 AG006786, R01 AG025711, R01 AG017216, R01 AG003949, NINDS grant R01 NS080820, CurePSP Foundation, and support from Mayo Foundation. Study data includes samples collected through the Sun Health Research Institute Brain and Body Donation Program of Sun City, Arizona. The Brain and Body Donation Program is supported by the National Institute of Neurological Disorders and Stroke (U24 NS072026 National Brain and Tissue Resource for Parkinsons Disease and Related Disorders), the National Institute on Aging (P30 AG19610 Arizona Alzheimers Disease Core Center), the Arizona Department of Health Services (contract 211002, Arizona Alzheimers Research Center), the Arizona Biomedical Research Commission (contracts 4001, 0011, 05-901 and 1001 to the Arizona Parkinson’s Disease Consortium) and the Michael J. Fox Foundation for Parkinsons Research.

We gratefully acknowledged Dr. Hyejung Won for his guidance to interpret the HiC sequencing data from the human fetal brains.

## DATA AVAILABILITY

All the data and analysis output are available via the AD Knowledge Portal (https://adknowledgeportal.synapse.org). The AD Knowledge Portal is a platform for accessing data, analyses, and tools generated by the Accelerating Medicines Partnership (AMP-AD) Target Discovery Program and other National Institute on Aging (NIA)-supported programs to enable open-science practices and accelerate translational learning. The data, analyses and tools are shared early in the research cycle without a publication embargo on secondary use. Data is available for general research use according to the following requirements for data access and data attribution (https://adknowledgeportal.synapse.org/DataAccess/Instructions). See data and supporting information: https://doi.org/10.7303/syn22240706

## REFERENCE

1. Farrer, L.A. et al. Effects of age, sex, and ethnicity on the association between apolipoprotein E genotype and Alzheimer disease. A meta-analysis. APOE and Alzheimer Disease Meta Analysis Consortium. JAMA 278, 1349–56 (1997).

2. Sherva, R. & Farrer, L.A. Power and pitfalls of the genome-wide association study approach to identify genes for Alzheimer’s disease. Curr Psychiatry Rep 13, 138–46 (2011).

3. Lambert, J.C. et al. Meta-analysis of 74,046 individuals identifies 11 new susceptibility loci for Alzheimer’s disease. Nat Genet 45, 1452–8 (2013).

4. Kunkle, B.W. et al. Genetic meta-analysis of diagnosed Alzheimer’s disease identifies new risk loci and implicates Abeta, tau, immunity and lipid processing. Nat Genet 51, 414–430 (2019).

5. Meyer, M.R. et al. APOE genotype predicts when--not whether--one is predisposed to develop Alzheimer disease. Nat Genet 19, 321–2 (1998).

6. Ma, Y. & Ordovas, J.M. The integration of epigenetics and genetics in nutrition research for CVD risk factors. Proc Nutr Soc 76, 333–346 (2017).

7. Ma, Y. et al. Analysis of Whole-Exome Sequencing Data for Alzheimer Disease Stratified by APOE Genotype. JAMA Neurol (2019).

8. De Jager, P.L. et al. Alzheimer’s disease: early alterations in brain DNA methylation at ANK1, BIN1, RHBDF2 and other loci. Nat Neurosci 17, 1156–63 (2014).

9. De Jager, P.L. et al. A multi-omic atlas of the human frontal cortex for aging and Alzheimer’s disease research. Sci Data 5, 180142 (2018).

10. Lunnon, K. et al. Methylomic profiling implicates cortical deregulation of ANK1 in Alzheimer’s disease. Nat Neurosci 17, 1164–70 (2014).

11. Smith, R.G. et al. Elevated DNA methylation across a 48-kb region spanning the HOXA gene cluster is associated with Alzheimer’s disease neuropathology. Alzheimers Dement 14, 1580–1588 (2018).

12. Zou, F. et al. Brain expression genome-wide association study (eGWAS) identifies human disease-associated variants. PLoS Genet 8, e1002707 (2012).

13. Allen, M. et al. Human whole genome genotype and transcriptome data for Alzheimer’s and other neurodegenerative diseases. Sci Data 3, 160089 (2016).

14. McKhann, G. et al. Clinical diagnosis of Alzheimer’s disease: report of the NINCDS-ADRDA Work Group under the auspices of Department of Health and Human Services Task Force on Alzheimer’s Disease. Neurology 34, 939–44 (1984).

15. Consensus recommendations for the postmortem diagnosis of Alzheimer’s disease. The National Institute on Aging, and Reagan Institute Working Group on Diagnostic Criteria for the Neuropathological Assessment of Alzheimer’s Disease. Neurobiol Aging 18, S1–2 (1997).

16. Bennett, D.A. et al. Neuropathology of older persons without cognitive impairment from two community-based studies. Neurology 66, 1837–44 (2006).

17. Yu, L., Boyle, P.A., Leurgans, S., Schneider, J.A. & Bennett, D.A. Disentangling the effects of age and APOE on neuropathology and late life cognitive decline. Neurobiol Aging 35, 819–26 (2014).

18. Bennett, D.A., Schneider, J.A., Arvanitakis, Z. & Wilson, R.S. Overview and findings from the religious orders study. Curr Alzheimer Res 9, 628–45 (2012).

19. Bennett, D.A. et al. Overview and findings from the rush Memory and Aging Project. Curr Alzheimer Res 9, 646–63 (2012).

20. White, C.C. et al. Identification of genes associated with dissociation of cognitive performance and neuropathological burden: Multistep analysis of genetic, epigenetic, and transcriptional data. PLoS Med 14, e1002287 (2017).

21. Pidsley, R. et al. A data-driven approach to preprocessing Illumina 450K methylation array data. BMC Genomics 14, 293 (2013).

22. Allen, M. et al. Gene expression, methylation and neuropathology correlations at progressive supranuclear palsy risk loci. Acta Neuropathol 132, 197–211 (2016).

23. Mostafavi, S. et al. A molecular network of the aging human brain provides insights into the pathology and cognitive decline of Alzheimer’s disease. Nat Neurosci 21, 811–819 (2018).

24. Felsky, D. et al. Neuropathological correlates and genetic architecture of microglial activation in elderly human brain. Nat Commun 10, 409 (2019).

25. Szklarczyk, D. et al. STRING v10: protein-protein interaction networks, integrated over the tree of life. Nucleic Acids Res 43, D447–52 (2015).

26. Benjamini Y H.Y. Controlling the false discovery rate: a practical and powerful approach to multiple testing. J. Roy. Statist. Soc. 57, 289–300 (1995).

27. Won, H. et al. Chromosome conformation elucidates regulatory relationships in developing human brain. Nature 538, 523–527 (2016).

28. Lesaffre, E.A., A. Partial separation in logistic discrimination. Journal of the Royal Statistical Society. Series B. Methodological 1, 109–116 (1989).

29. Patrick, E.T. M.; Ergun, A.; Ng, B.; Casazza, W.; Cimpean, M.; Yung, C.; Schneider, J.A.; Bennett, D.A.; Gaiteri, C.; De Jager P.L.; Bradshaw, E.M.; Mostafavi, S. Deconvolving the contributions of cell-type heterogeneity on cortical gene expression. bioRxiv (2019).

30. Jun, G. et al. A novel Alzheimer disease locus located near the gene encoding tau protein. Mol Psychiatry 21, 108–17 (2016).

31. Strickland, S.L. et al. MAPT haplotype-stratified GWAS reveals differential association for AD risk variants. Alzheimers Dement (2020).

