## Supplementary File for "Epigenomic features related to microglia are associated with attenuated effect of APOE ε4 on alzheimer’s disease risk in humans"

**SUPPLEMENTAL MATERIAL CONTENTS**

**SUPPLEMENTAL METHODS (Text)**

**SUPPLEMENTAL TABLES**

**Table S1.** Population demographics included in each analysis across all studies.

**Table S2.** Summary statistics of the 25 CpG dinucleotides for their associations with the 11 common neuropathologies in all subjects of ROSMAP, and meta-interactions with APOE ε4 on pathological AD in all subjects across ROSMAP, LBB, and MSBB, and meta-associations with pathological AD in subgroups of subjects carrying or not carrying ε4 allele across ROSMAP, LBB, and MSBB (in excel spreadsheet).

**Table S3.** Meta-analysis of cg08706567 and cg26884773 using penalized generalized regression model.

**Table S4.** Interaction and association tests with scaled continuous variables.

**Table S5.** Summary statistics of the top 71 genes from the transcriptome wide association for the PC1 and cg12370200 in ROSMAP with the adjustment of *APOE* ε4 status MSBB (in excel spreadsheet).

**Table S6.** Summary statistics of the association between the 47 gene modules and the PC1 and cg12370200 in ROSMAP with the adjustment of *APOE* ε4 status MSBB (in excel spreadsheet).

**Table S7.** Associations of PC1 and cg12307200 with microglia cell type proportion in subset of subjects with immuno-histochemistry measurements (N=57).

**SUPPLEMENTAL FIGURES**

**Figure S1.** Forest plots of the association between the pathological diagnosis of Alzheimer’s disease (AD) and methylation level at the 4 top CpG dinucleotides.

**Figure S2.** Distributions of the scaled values of each of the top 4 CpG dinucleotides within each cohort before the derivation of PC1 and their pairwise correlations.

**Figure S3.** Associations of the proportion of activated microglia (PAM) with the methylation PC1 and cg12307200.

**SUPPLEMENTAL METHODS**

*Study description of ROSMAP, LBB, and MSBB*

The Religious order Study (ROS) and the Memory and Aging Project (MAP) is a longitudinal community-based study which already has recruited >3,000 dementia-free subjects at baseline. The participants have detailed cognitive, neuroimaging and other ante-mortem phenotyping and structured neuropathologic examinations at the time of death and postmortem multi-omic measurements. We also obtained DNA methylation data of the brain tissues archived in the MRC London Neurodegenerative Disease Brain Bank (LBB) (http://www.kcl.ac.uk/iop/depts/cn/research/MRC-London-Neurodegenerative-Diseases-Brain-Bank/MRC-London-Neurodegenerative-Diseases-Brain-Bank.aspx). Ethical approval for the study was provided by the NHS South East London REC 3. We also included DNA methylation data obtained from the brain tissues stored in the Mount Sinai Alzheimer’s Disease and Schizophrenia Brain Bank (MSBB) (<http://icahn.mssm.edu/researach/labs/neuropathology-and-brain-banking>).

*Neuropathologic indices in ROSMAP*

Details of the 11 common neuropathologic indices in ROSMAP were described before [1-4]. In brief, neuronal neurofibrillary tangles are identified by molecularly specific immunohistochemistry (antibodies to abnormally phosphorylated Tau protein, AT8). The mean of values in 8 regions were used into the analysis, which are hippocampus, entorhinal cortex, midfrontal cortex, inferior temporal, angular gyrus, calcarine cortex, anterior cingulate cortex, and superior frontal cortex [2, 3]. Neuritic plaques and diffuse plaques are the means of the standard deviation scaled regional counts of the microscopic silver-stained slides from five regions: midfrontal cortex, midtemporal cortex, inferior parietal cortex, entorhinal cortex, and hippocampus. Lewy body pathology is defined as a binary variable (presence or not) using immunohistochemistry in nigra and/or cortex [5, 6]. Macroscopic cerebral infarctions are defined as presence of one or more gross chronic cerebral infarctions visible to the naked eye on fixed slabs which are dissected for histological confirmation [7]. Microscopic cerebral infarctions are defined as presence of one or more chronic microinfarcts which are examined a minimum of nine regions in one hemisphere on 6 μm paraffin-embedded sections with H&E stain[8]. A four-level severity scale was applied to measure the atherosclerosis and arteriosclerosis according to the large and small vessel pathologies in anterior basal ganglia [9]. Cerebral amyloid angiopathy (CAA) was graded on a five-level scale (0 to 4) in four neocortical regions (mid-frontal, angular gyrus, inferior temporal gyrus, and calcarine cortex) and averaged to derive a CAA score, as previously described [10]. Hippocampal sclerosis was recorded as either present or absent as evaluated with H&E stain as described before [4]. Transactive response DNA-binding protein 43 kDa (TDP-43) proteinopathy was measured and categorized into four levels of severity as previously described [11]: no inclusions (stage 0), inclusions in amygdala only (stage 1), inclusions in amygdala as well as entorhinal cortex and/or hippocampus CA1 (stage 2), and inclusions in amygdala, neocortex, and entorhinal cortex and/or hippocampus CA1 (stage 3).

*Brain gene expression in ROSMAP, MSBB, and MAYO*

ROSMAP: As reported previously[12], RNA extracted from the gray matter of ROSMAP cortical samples were followed with library preparation using the strand specific dUTP protocol with poly-A selection, Illumina HiSeq sequencing (101bp paired-end reads with 50M reads coverage), STAR[13] alignment, RSEM v1.2.31[14] expression estimates, and QC to remove genes with <15 median expected count of raw expression or having outlying mean expression or variance and subjects with poor quality data or duplicated samples. The gene expression levels were log2 transformed and were corrected using TMM normalization factors using voom of the “edgeR”[15], followed with linear corrections for batch, median 3’ bias (correlated with RIN), and number of ribosomal bases using “limma”. The final output values were the normalized residuals on the log2(cpm) scale of the 17,068 autosomal genes across 421 subjects. A subset of these subjects (N=413) were previously used to derive the 47 cell-type relevant gene module[16].

MSBB: With the genotype concordance check based on the 58 single nucleotide polymorphisms included in both the DNA methylation data and the whole genome sequencing data, we have identified 50 subjects (concordance rate >0.90) who have been profiled with both DNA methylation at prefrontal cortex and RNA-seq at BM44 region (closest to the prefrontal cortex). We downloaded the gene expressions of the transcriptome from Synapse platform ([https://www.synapse.org/#!Synapse:syn7391](https://www.synapse.org/#!Synapse:syn8691099.1)833). The detailed descriptions of the methods were described online (<https://www.synapse.org/#!Synapse:syn17010685)>. In brief, the gene expression values are represented by the counts of the aligned reads regarding to each gene based on the reference of GENCODE24 (GRCh38) using STAR by setting “quantMode” as “GeneCounts”.

*MAYO*: We included 45 AD cases with both the DNA methylation data at cg05157625 and gene expression data from their temporal cortex. As described before, the gene expression profiles were measured by the Illumina Whole Genome DASL (WG-DASL) microarray[17, 18].

*Statistical analysis*

During the post-hoc check, we found that for the cg08706567 and cg26884773, the LBB ε4+ dataset did not contribute to the meta-analysis since the dataset encounter a problem of complete separation and infinite maximum likelihood estimates when the logistic model included all the covariates of age at death, sex, postmortem interval, and cell proportion. The modeling yielded the warning message of "glm.fit: fitted probabilities numerically 0 or 1 occurred" and provided the inflated beta coefficient, standard error of the predictors (cg08706567 and cg26884773) and *P* values of 1 as observed in the presented forest plot. We have conducted the separation detection using the R package of "detectseparation" and confirmed that the combination of all the covariates and these two CpG dinucleotides in LBB ε4+ dataset leads to this problem. The reference of Lesaffre, E., and Albert, A. 1989 [19] described the situation when these infinite estimates might happen. The model is successively refitted by increasing the maximum number of allowed iteratively reweighted least squares iterations at each step, and the asymptotically estimated standard errors from each step are divided to the corresponding ones from the first fit. If the sequence of ratios diverges, then the maximum likelihood estimate is minus or plus infinity, which were observed in our cases. One approach to bypass this problem is to run the penalized maximum likelihood methods with a quasi Fisher scoring iteration implemented in the R package of "brglm2" (https://cran.r-project.org/web/packages/brglm2/index.html). We presented the results in Table S5 where meta-P values with both LBB and MSBB was 0.01 for cg08706567 and 7.35E-03 for cg26884773. In this case, all the 4 top CpG dinucleotides we have identified have nominal significance during the meta-analysis within only LBB and MSBB. During our post-hoc check of the LBB ε4+ dataset, there are 9 out of the 25 tested CpG dinucleotides with P≥0.99 (an indication of the separation problem as cg08706567 and cg26884773), however none of the CpG dinucleotides have P≥0.99 in the LBB all subjects (ε4+ and ε4-). Considering the fact that those penalized methods are not idea to provide the exactly accurate estimates, we still rely on the un-penalized method for the discovery of top CpG dinucleotides.

**Table S1.** Population demographics included in each analysis across all studies.

|  | ROSMAP | | | | | |  | LBB |  | MSBB | |  | MAYO |
| --- | --- | --- | --- | --- | --- | --- | --- | --- | --- | --- | --- | --- | --- |
|  | MWAS | TWAS | modWAS | IHC | | |  | Methylation analysis |  | Methylation analysis | TWAS replication |  | TWAS replication |
|  |  |  |  | All with PAM measurement | With both PAM and DNA methylation data | With different cell type proportion measurement and DNA methylation data |  |  |  |  |  |  |  |
| All subjects |  |  |  |  |  |  |  |  |  |  |  |  |  |
| N | 572 | 421 | 413 | 136 | 122 | 57 |  | 68 |  | 129 | 50 |  | 45 |
| Age at death (y) | 88.32 ± 6.50 | 88.61 ± 6.66 | 88.63 ± 6.61 | 89.24 ± 5.08 | 89.57 ± 4.88 | 85.68 ± 6.73 |  | 84.65 ± 9.48 |  | 85.63 ± 7.60 | 85.42 ± 7.82 |  | 73.82 ± 5.61 |
| Female (N, %) | 361, 63.11% | 256, 62.95% | 259, 62.71% | 88, 64.71% | 78, 63.93% | 0, 0% |  | 46, 67.65% |  | 49, 62.02% | 29, 58% |  | 19, 42.2% |
| ε4+ (N, %) | 158, 27.62% | 111, 26.37% | 109, 26.39% | 37, 27.21% | 32, 26.23% | 16, 28.07% |  | 36, 52.94% |  | 41, 31.78% | 17, 34% |  | 27, 60% |
| Pathological AD (N, %) | 349, 61.01% | 250, 59.38% | 243, 58.84% | 82, 60.29% | 74, 60.66% | 29, 50.88% |  | 52, 76.47% |  | 74, 57.36% | 29, 58% |  | 45, 100% |
| ε4+ subgroup |  |  |  |  |  |  |  |  |  |  |  |  |  |
| N | 158 | 111 | 109 | 37 | 32 | 16 |  | 36 |  | 41 | 17 |  | 27 |
| Age at death (y) | 87.31 ± 6.01 | 88.04 ± 5.91 | 88.08 ± 5.89 | 87.81 ± 5.01 | 88.27 ± 4.58 | 87.39 ± 7.87 |  | 84.58 ± 9.00 |  | 86.27 ± 7.22 | 86.35 ± 8.91 |  | 73.04 ± 6.12 |
| Female (N, %) | 93, 58.86% | 46, 58.56% | 63, 57.80% | 23, 62.16% | 20, 62.5% | 0, 0% |  | 25, 69% |  | 27, 66% | 10, 58.82% |  | 11, 40.74% |
| Pathological AD (N, %) | 128, 81.01% | 89, 80.18% | 87, 79.82% | 30, 81.08% | 26, 81.25% | 12, 75% |  | 33, 92% |  | 34 83% | 14, 82.35% |  | 27, 100% |
| ε4- subgroup |  |  |  |  |  |  |  |  |  |  |  |  |  |
| N | 414 | 310 | 304 | 99 | 90 | 41 |  | 32 |  | 88 | 33 |  | 18 |
| Age at death (y) | 88.70 ± 6.65 | 88.82 ± 6.91 | 88.82 ± 6.85 | 89.78 ± 5.03 | 90.04 ± 4.92 | 85.01 ± 6.21 |  | 84.72 ± 10.13 |  | 85.33 ± 7.80 | 84.94 ± 7.30 |  | 75 ± 4.64 |
| Female (N, %) | 268, 64.73% | 200, 64.52% | 196, 64.47% | 65, 65.66% | 58, 64.44% | 0, 0% |  | 21, 66% |  | 53, 60% | 19, 57.58% |  | 8, 44.44% |
| Pathological AD (N, %) | 221, 53.38% | 161, 51.94% | 156, 51.32% | 52, 52.53% | 48, 53.33% | 17, 41.46% |  | 19, 59% |  | 40, 45% | 15, 45.45% |  | 18, 100% |

Abbreviations: MWAS, methylome-wide association analysis; TWAS, transcriptome-wide association analysis; modWAS, co-expression modules genomewide association analysis; IHC, immuno-histochemistry measurements.

**Table S3.** Meta-analysis of cg08706567 and cg26884773 using penalized generalized regression model.

|  | cg08706567 | | | |  | cg26884773 | | | |
| --- | --- | --- | --- | --- | --- | --- | --- | --- | --- |
|  | N | BETA | STDERR | *P* |  | N | BETA | STDERR | *P* |
| ROSMAP | 158 | 34.30 | 9.52 | 3.14E-04 |  | 158 | 33.70 | 9.26 | 2.76E-04 |
| *LBB | 36 | 54.10 | 27.26 | 0.05 |  | 36 | 42.21 | 22.33 | 0.06 |
| *MSBB | 41 | 27.50 | 16.20 | 0.09 |  | 41 | 44.90 | 23.60 | 0.06 |
| *META | 77 | 34.44 | 13.93 | 0.01 |  | 77 | 43.48 | 16.22 | 7.35E-03 |
| META_All | 235 | 34.35 | 7.86 | 1.24E-05 |  | 235 | 36.10 | 8.04 | 7.14E-06 |

Note: BETA, STDERR, and *P* represent the regression coefficient, standard error and corresponding *P* value of the effect of the methylation of the CpG dinucleotide on the risk of AD.

In the LBB dataset, the penalized regression model was conducted. *META means the meta-analysis between the LBB and MSBB.

**Table S4.** Interaction and association tests with scaled continuous variables.

| Variable |  | N | BETA^1^ | STDERR^1^ | *P^1^* | *P^2^* |
| --- | --- | --- | --- | --- | --- | --- |
| PC1 | Ιnteraction with ε4 | 767 | 0.98 | 0.29 | 7.26E-04 | 7.21E-04 |
|  | Main effect in ε4+ | 235 | 1.36 | 0.30 | 7.58E-06 | 7.40E-06 |
|  | Main effect in ε4- | 532 | 0.32 | 0.11 | 2.59E-03 | 2.66E-03 |
| cg12307200 | Ιnteraction with ε4 | 767 | -0.50 | 0.27 | 5.80E-02 | 6.48E-02 |
|  | Main effect in ε4+ | 235 | -1.15 | 0.28 | 4.67E-05 | 5.74E-05 |
|  | Main effect in ε4- | 532 | -0.50 | 0.11 | 9.99E-06 | 7.93E-06 |

^1^BETA, STDERR and *P* represent the regression coefficient and its corresponding standard error and *P* value for the effect of the scaled values of each of the listed variable on the risk of pathological AD in the logistic regression model with the adjustment of sex, methylation experimental batches, study, and scaled values of all the continuous covariates including age at death, postmortem interval and two major ethnicity principles.

^2^*P* represent the *P* value of the same variable which is not scaled and all the continuous covariates are not scaled either.

**Table S7.** Associations of PC1 and cg12307200 with microglia cell type proportion in subset of subjects with immuno-histochemistry measurements (N=57).

| Exposure variable | Approach | Measurement | Outcome variable | BETA | STDERR | *P* |
| --- | --- | --- | --- | --- | --- | --- |
| PC1 | IHC | IBA1 protein | Microglia cell type proportion | 0.005 | 3.78 | 0.23 |
|  | RNA-seq | *IBA1* mRNA | IBA-1 mRNA expression | -0.03 | 0.09 | 0.78 |
| cg12307200 | IHC | IBA1 protein | Microglia cell type proportion | -0.15 | 0.11 | 0.17 |
|  | RNA-seq | *IBA1* mRNA | IBA-1 mRNA expression | -0.53 | 2.63 | 0.84 |

Note: BETA, STDERR, and *P* represent the regression coefficient, standard error and corresponding *P* value of the effect of the exposure variable on the outcome variable using the general linear regression model. The outcome variable of the IHC approach is the proportion of microglia cells measured by IBA-1 protein staining out of the total number of cells measured, including neurons, astrocytes, oligodendrocytes, endothelial cells, and microglia. The covariates include *APOE* ε4 carrying status, age at death, postmortem interval, methylation experimental batch and two major ethnic principal components. Abbreviation: IHC, immuno-histochemistry.

**Figure S1. Forest plots of the association between the pathological diagnosis of Alzheimer’s disease (AD) and methylation level at the 4 top CpG dinucleotides.** The results were shown for (A) cg12307200, (B) cg05157625, (C) cg08706567, and (D) cg26884773. The association were conducted within: 1) all the subjects; 2) subjects carrying; or 3) not carrying the *APOE* ε4 allele; and 4) the interaction test between the methylation level and *APOE* ε4 allele carrying status within all the subjects. The filled square and horizontal line for each population or the filled summary diamonds (blue for the meta-analysis across the replication cohorts while red for the joint meta-analysis across all the cohorts) denote the estimated regression coefficient (BETA) and its 95% CI per unit increase in the methylation level of each CpG dinucleotide or the interaction term of the methylation times the ε4-carrying status (yes=1 and no=0) for the binary outcome variable of the pathological diagnosis of AD (no=0 and yes=1) with the adjustment of the covariates of age at death, postmortem interval, sex, study, ethnicity principle components, methylation experiment batches in ROSMAP, and cell proportion in LBB and MSBB.

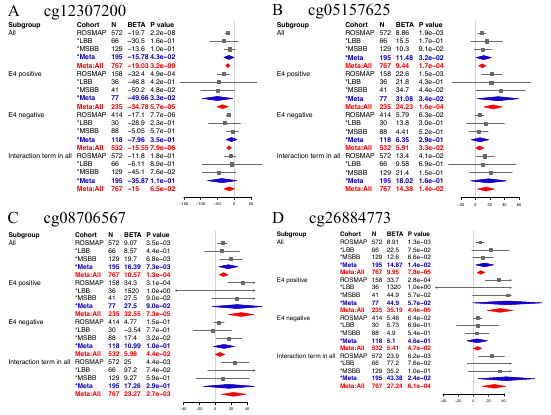

**Figure S2. Distributions of the scaled values of each of the top 4 CpG dinucleotides within each cohort before the derivation of PC1 and their pairwise correlations.** (A-C) Histogram shows the distribution of the scaled β values of each CpG dinucleotide (pink for cg08706567, yellow for cg26884773, blue for cg12307200, and red for cg05157625) within ROSMAP (A), LBB (B), and MSBB (C). (D-F) Correlation matrix among the top 4 CpGs (unscaled values) within ROSMAP (D), LBB (E), and MSBB (F). The correlation coefficients are presented and color-coded with red for positive correlation and blue for the negative correlation.

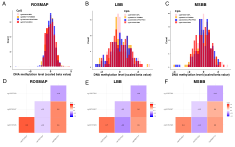

**Figure S3. Associations of the proportion of activated microglia (PAM) with the methylation PC1 and cg12307200.** The *P* values with the sign of the effect direction of the association results for PAM at inferior temporal cortex (horizontal axis), midfrontal cortex (right diagonal axis), posterior putamen (vertical axis), and ventral medial caudate (left diagonal axis). The purple dashed circle represents the nominal significance of *P* = 0.05. The sign and *P* values were obtained through the generalized linear regression model, in which the outcome variable is the PAM (square root), the exposure variable is the methylation PC1 (red dot) or cg12307200 (blue dot), and the covariates include age at death, sex, postmortem interval, study, *APOE* ε4 binary status, two major ethnic principles and methylation experimental batches.

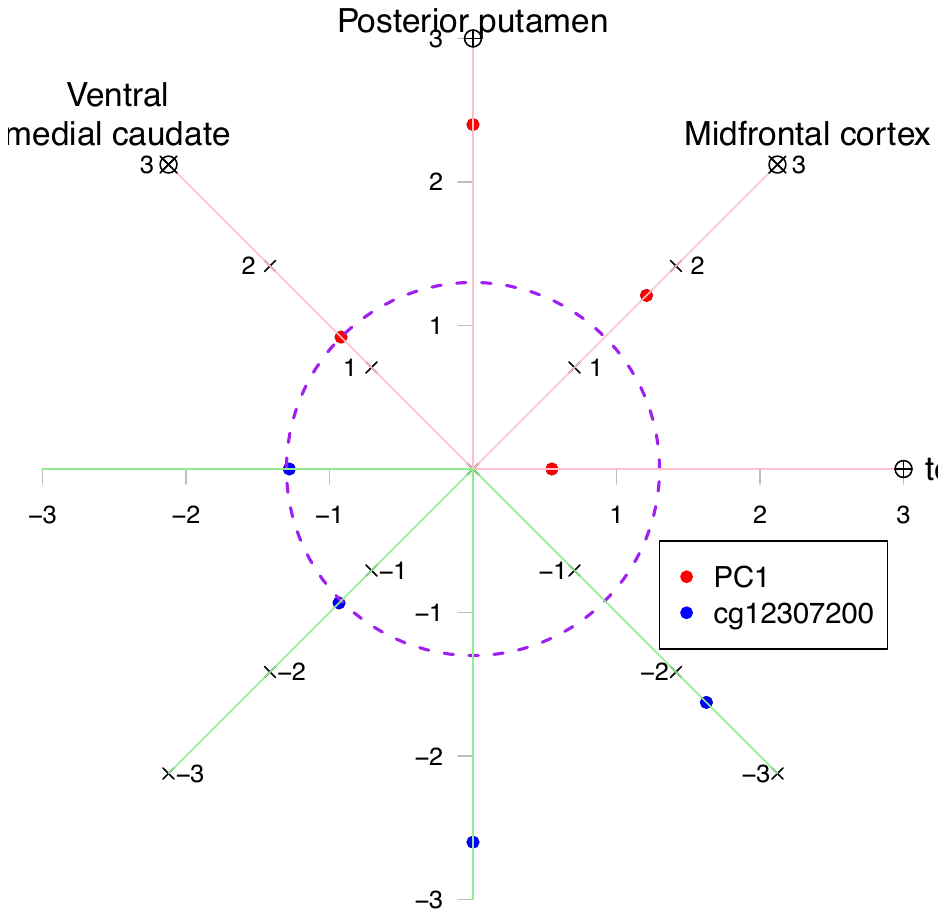

**References:**

1.	Yu L, Boyle PA, Leurgans S, Schneider JA, Bennett DA. Disentangling the effects of age and APOE on neuropathology and late life cognitive decline. Neurobiol Aging. 2014;35(4):819-26. doi: 10.1016/j.neurobiolaging.2013.10.074. PubMed PMID: 24199961; PubMed Central PMCID: PMCPMC3882071.

2.	Bennett DA, Schneider JA, Arvanitakis Z, Wilson RS. Overview and findings from the religious orders study. Curr Alzheimer Res. 2012;9(6):628-45. PubMed PMID: 22471860; PubMed Central PMCID: PMCPMC3409291.

3.	Bennett DA, Schneider JA, Buchman AS, Barnes LL, Boyle PA, Wilson RS. Overview and findings from the rush Memory and Aging Project. Curr Alzheimer Res. 2012;9(6):646-63. PubMed PMID: 22471867; PubMed Central PMCID: PMCPMC3439198.

4.	White CC, Yang HS, Yu L, Chibnik LB, Dawe RJ, Yang J, et al. Identification of genes associated with dissociation of cognitive performance and neuropathological burden: Multistep analysis of genetic, epigenetic, and transcriptional data. PLoS Med. 2017;14(4):e1002287. doi: 10.1371/journal.pmed.1002287. PubMed PMID: 28441426; PubMed Central PMCID: PMCPMC5404753.

5.	Buchman AS, Boyle PA, Wilson RS, Tang Y, Bennett DA. Frailty is associated with incident Alzheimer's disease and cognitive decline in the elderly. Psychosom Med. 2007;69(5):483-9. doi: 10.1097/psy.0b013e318068de1d. PubMed PMID: 17556640.

6.	Wilson RS, Yu L, Schneider JA, Arnold SE, Buchman AS, Bennett DA. Lewy bodies and olfactory dysfunction in old age. Chem Senses. 2011;36(4):367-73. doi: 10.1093/chemse/bjq139. PubMed PMID: 21257733; PubMed Central PMCID: PMCPMC3073534.

7.	Schneider JA, Bienias JL, Wilson RS, Berry-Kravis E, Evans DA, Bennett DA. The apolipoprotein E epsilon4 allele increases the odds of chronic cerebral infarction [corrected] detected at autopsy in older persons. Stroke. 2005;36(5):954-9. doi: 10.1161/01.STR.0000160747.27470.2a. PubMed PMID: 15774818.

8.	Arvanitakis Z, Leurgans SE, Barnes LL, Bennett DA, Schneider JA. Microinfarct pathology, dementia, and cognitive systems. Stroke. 2011;42(3):722-7. doi: 10.1161/STROKEAHA.110.595082. PubMed PMID: 21212395; PubMed Central PMCID: PMCPMC3042494.

9.	Yip AG, McKee AC, Green RC, Wells J, Young H, Cupples LA, et al. APOE, vascular pathology, and the AD brain. Neurology. 2005;65(2):259-65. doi: 10.1212/01.wnl.0000168863.49053.4d. PubMed PMID: 16043796.

10.	Boyle PA, Yu L, Nag S, Leurgans S, Wilson RS, Bennett DA, et al. Cerebral amyloid angiopathy and cognitive outcomes in community-based older persons. Neurology. 2015;85(22):1930-6. doi: 10.1212/WNL.0000000000002175. PubMed PMID: 26537052; PubMed Central PMCID: PMCPMC4664125.

11.	Yu L, De Jager PL, Yang J, Trojanowski JQ, Bennett DA, Schneider JA. The TMEM106B locus and TDP-43 pathology in older persons without FTLD. Neurology. 2015;84(9):927-34. doi: 10.1212/WNL.0000000000001313. PubMed PMID: 25653292; PubMed Central PMCID: PMCPMC4351662.

12.	De Jager PL, Ma Y, McCabe C, Xu J, Vardarajan BN, Felsky D, et al. A multi-omic atlas of the human frontal cortex for aging and Alzheimer's disease research. Sci Data. 2018;5:180142. doi: 10.1038/sdata.2018.142. PubMed PMID: 30084846; PubMed Central PMCID: PMCPMC6080491.

13.	Dobin A, Davis CA, Schlesinger F, Drenkow J, Zaleski C, Jha S, et al. STAR: ultrafast universal RNA-seq aligner. Bioinformatics. 2013;29(1):15-21. doi: 10.1093/bioinformatics/bts635. PubMed PMID: 23104886; PubMed Central PMCID: PMCPMC3530905.

14.	Li B, Dewey CN. RSEM: accurate transcript quantification from RNA-Seq data with or without a reference genome. BMC Bioinformatics. 2011;12:323. doi: 10.1186/1471-2105-12-323. PubMed PMID: 21816040; PubMed Central PMCID: PMCPMC3163565.

15.	Robinson MD, McCarthy DJ, Smyth GK. edgeR: a Bioconductor package for differential expression analysis of digital gene expression data. Bioinformatics. 2010;26(1):139-40. doi: 10.1093/bioinformatics/btp616. PubMed PMID: 19910308; PubMed Central PMCID: PMCPMC2796818.

16.	Mostafavi S, Gaiteri C, Sullivan SE, White CC, Tasaki S, Xu J, et al. A molecular network of the aging human brain provides insights into the pathology and cognitive decline of Alzheimer's disease. Nat Neurosci. 2018;21(6):811-9. doi: 10.1038/s41593-018-0154-9. PubMed PMID: 29802388.

17.	Zou F, Chai HS, Younkin CS, Allen M, Crook J, Pankratz VS, et al. Brain expression genome-wide association study (eGWAS) identifies human disease-associated variants. PLoS Genet. 2012;8(6):e1002707. doi: 10.1371/journal.pgen.1002707. PubMed PMID: 22685416; PubMed Central PMCID: PMCPMC3369937 from Elan Pharmaceutical Research, Pfizer Pharmaceuticals, Medivation, and Forrest. RC Petersen has been a consultant to GE Healthcare and Elan Pharmaceuticals, has served on a data safety monitoring committee for Pfizer and Janssen Alzheimer Immunotherapy, and has provided a CME lecture for Novartis. ADGC Authors: T.D.B. received licensing fees from and is on the speaker's bureau of Athena Diagnostics. M.R.F. receives research funding from BristolMyersSquibb Company, Danone Research, Elan Pharmaceuticals, Eli Lilly, Novartis Pharmaceuticals, OctaPharma AG, Pfizer, and Sonexa Therapeutics; receives honoraria as scientific consultant from Accera, Astellas Pharma US Inc., Baxter, Bayer Pharmaceuticals Corporation, BristolMyersSquibb, Eisai Medical Research, GE Healthcare, Medavante, Medivation, Merck, Novartis Pharmaceuticals, Pfizer, Prana Biotechnology, QR Pharma, The sanofi-aventis Group, and Toyama Chemical; and is speaker for Eisai Medical Research, Forest Laboratories, Pfizer, and Novartis Pharmaceuticals. A.M.G. has research funding from AstraZeneca, Pfizer and Genentech, and has received remuneration for giving talks at Pfizer and Genentech. R.C.P. is on the Safety Monitory Committee of Pfizer (Wyeth) and a consultant to the Safety Monitoring Committee at Janssen Alzheimer's Immunotherapy Program (Elan), to Elan Pharmaceuticals, and to GE Healthcare. R.E.T. is a consultant to Eisai, Japan, in the area of Alzheimer's genetics and a shareholder in, and consultant to, Pathway Genomics, San Diego, CA.

18.	Allen M, Carrasquillo MM, Funk C, Heavner BD, Zou F, Younkin CS, et al. Human whole genome genotype and transcriptome data for Alzheimer's and other neurodegenerative diseases. Sci Data. 2016;3:160089. doi: 10.1038/sdata.2016.89. PubMed PMID: 27727239; PubMed Central PMCID: PMCPMC5058336 Immunotherapy: Chair, Data Monitoring Committee. Hoffman-La Roche, Inc.: Consultant. Merck, Inc.: Consultant. Genentech, Inc.: Consultant. Biogen, Inc.: Consultant. Eli Lilly & Co.: Consultant. N.R.G.-R. has multicenter treatment study grants from Lilly and TauRx and consulted for Cytox. N.E.-T. has consulted for Cytox. The remaining authors declare no competing financial interests.

19.	Lesaffre EA, A. Partial separation in logistic discrimination. Journal of the Royal Statistical Society Series B Methodological. 1989;1(51):109-16. doi: https://doi.org/10.1111/j.2517-6161.1989.tb01752.x.
